# Beyond accuracy, precision and TAE: Direct assay validation against product specification aligned with USP <1033>

**DOI:** 10.1101/2025.05.21.655192

**Authors:** Davor Josipovic

**Affiliations:** Independent

**Keywords:** USP <1033>, ICH Q2(R2), analytical procedure validation, accuracy and precision, total analytical error, bioassay validation, assay validation, potency assay validation, analytical target profile, validation methodology

## Abstract

Current validation methodologies – whether based on accuracy and precision or total analytical error (TAE) and risk – fall short when the analytical target profile (ATP) is defined in terms of product and process specifications, as is the case in the USP <1033> guideline for biological assay validation. In this paper, we critically examine the limitations of these existing methodologies and introduce a statistically correct methodology tailored to the ATP formulation. While this novel methodology is demonstrated in the context of potency assay validation in line with the USP <1033> guideline, it is broadly applicable to other analytical procedures governed by ICH Q2(R2), with only minor adaptations. A freely accessible online application has also been developed to facilitate discussion and adoption of the novel methodology in practice.

## Introduction

The principal guideline for validation of analytical procedures within the pharmaceutical industry is ICH Q2(R2) [1] which outlines two major methodologies for validation of analytical procedures: one based on accuracy and precision, and other based on total analytical error (TAE) and risk.

The official USP <1033> [2] guideline follows the accuracy and precision methodology, while its latest draft [3] also introduces the TAE methodology, applying both to bioassays, in particular the potency assays. Before these validation methodologies can be applied however, validation acceptance criteria need to be defined. USP <1033> [2] states that “ when there is an existing product specification, acceptance criteria can be justified on the basis of the risk that measurements may fall outside of the product specification”. This requirement is usually formalized in the Analytical Target Profile (ATP) of the procedure. Then the guideline proposes how to derive acceptance criteria for accuracy and precision or TAE and risk, such that they comply with the ATP.

In this paper we show that both of these methodologies fall short in capturing the essence of the ATP when it is defined in terms of product specifications, and that to correctly validate the procedure a novel validation methodology is required.

The paper is structured as follows. In the first section the general assumptions are made explicit in a measurement model that serves as the foundation for later inference. In subsequent section the ATP is defined and formalized based on the measurement model. In the next section accuracy and precision methodology is assessed, showing how it fails to meet the ATP requirements. A similar conclusion is drawn in the section on TAE-based validation. Then the correct validation methodology is introduced, and it is shown how it fully satisfies the ATP requirements and how it resolves the shortcomings of the prior two methodologies. The final discussion section addresses important but peripheral topics not covered in the main body of the paper.

### The assumptions

To properly address the research question, it is essential to begin with clear definitions and explicit assumptions. Central to this is the measurement model, which formally describes how the analytical procedure measures relative potency (RP).

#### The measurement model

A measurement model that is consistent with USP <1033> [2] definitions and formulas is the following:

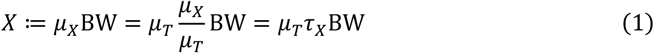

where *X* is the measured value, *μ*_*T*_ the true (target) relative potency, *τ*_*X*_ = *μ*_*X*_/*μ*_*T*_ the trueness factor (or multiplicative systematic error), and 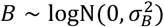 and 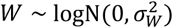 the independent lognormal random variables with 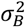 the between and 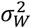 the within run variability of the procedure.

Concretely, this means that every time one measures relative potency, the measured value *X* can be decomposed into the true potency value of the measurand, the systematic bias and the random realizations of between and within variance components of the procedure. Note that the terms “ measurement”, measured “ value”, “ measurand”, “ true value” and “ systematic error” are borrowed from GUM [4] in their original definitions.

Furthermore, knowing that the product of two independent lognormal random variables is lognormal, one can also deduce that:

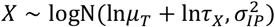

where for convenience 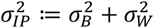.

Lastly, for plotting purposes, *X* is often represented as relative error (%) through the following transformation:

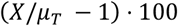

#### Measuring production samples

When the relative potency of a manufactured product is measured with this procedure, then the true value (*μ*_*T*_) in the measurement model is the true (albeit unknown) relative potency of that manufactured product. Assuming the production process produces products with 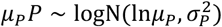 relative potency, we can substitute *μ*_*T*_ with *μ*_*p*_*P* in Eq. 1. Our measured value *X* becomes then the measured value of one realization of this production process:

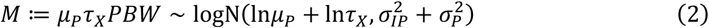

Now 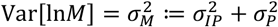, which means that the observed variability of the logarithm of our measured potency value is composed of the production process variability and analytical procedure variability, just as stated in the latest USP <1033> draft [3], but now deduced formally from a measurement model which can be used for inference.

The measured value refers to the output of a single implementation of the procedure on a test article, ideally from one run and one replicate per run. In contrast, the reportable value (*RV*) [5] extends this concept by being a function of a set of measured values, hence *RV* ≔ *f*({*M*_*nk*_}) where *n* is the run and *k* the replicate index. The function *f* is usually – but not necessarily – the geometric mean.

#### Global versus local performance parameters

Lastly make note of an important distinction between the parameters *τ*_*X*_ and 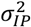, and *τ*_*X*_(*μ*_*T*_) and 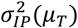 respectively. The former are called global because they are fixed (i.e. constant) scalars, and the later are called local because they are defined as (unknown) functions depending on the true value. Local parameters are more general and can be freely substituted in place of the global ones in the measurement model above. The distinction between the two is important because during validation these parameters are estimated in function of the true value (and thus *depend* on the true value, cf. Table I) and are local, while during acceptance criteria determination in the USP <1033> [2,3] guideline, global parameters are used. This simplification results in excessively strict acceptance criteria and will be discussed in due time.

**Table I:** The locality of the procedure’s performance. At each “ local” level, the performance of the procedure is expressed in terms of τ_X_ and σ_IP_, or its derivatives RB and IP. The values are taken directly from the USP <1033> [2] example.

| Level (RP) | $\tau_X$ (RP) | $RB$ (%) | $\sigma_{IP}$ (RP) | $IP_{GCV}$ (%) |
| --- | --- | --- | --- | --- |
| **0.50** | 1.0323 | 3.23 | 0.0658 | 6.81 |
| **0.71** | 1.0006 | 0.06 | 0.0704 | 7.29 |
| **1.00** | 1.0497 | 4.97 | 0.0812 | 8.46 |
| **1.41** | 1.0291 | 2.91 | 0.0609 | 6.28 |
| **2.00** | 1.0972 | 9.72 | 0.0699 | 7.24 |

### Analytical Target Profile (ATP)

Before initiating validation, an Analytical Target Profile (ATP) must be defined. According to ICH Q14 [6] the ATP is a statement that defines “ the performance characteristics describing the intended purpose and the anticipated performance criteria of an analytical measurement”.

In USP <1033> [2] the ATP is defined in relation to product (release) specification (e.g. [0.71; 1.41] RP), a potential manufacturing bias, and the requirement that whenever the relative potency of a manufactured product is reported, the probability that this reportable value is out of product specification is not more than 1 %. The guideline also states that “ product variability is assumed to be equal to 0 in the calculations.”

The latest USP <1033> draft [3] builds on these assumptions but introduces a more realistic model for the production process. Instead of assuming zero variability, the draft assumes that the distribution of the potency of the manufactured product follows a lognormal distribution with a defined geometric mean (*μ*_*P*_) and geometric standard deviation (*σ*_*P*_). This is consistent with our previous definition of a production process *μ*_*P*_*P*.

While production process variability cannot be directly controlled, one can define validation criteria for the analytical procedure to ensure that failures due to measurement variability remain acceptably low and within product specification. Hence the above requirement results in the following ATP example where for the sake of simplicity we base it on the most elementary form of the reportable value:

> *[ATP] The procedure must be able to quantify relative potency (RP) in a range from 0*.*5 to 2*.*0 RP such that, under an assumed lognormal manufacturing distribution geometrically centered on 1*.*0 RP with a geometric standard deviation of 0*.*0477 RP, the expected probability of reportable values (one run, one replicate) falling outside [0*.*70; 1*.*43] RP is less than 1%*.

The general form of this ATP can be formalized as:

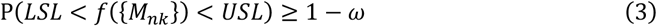

where *LSL* and *USL* denote the lower and upper product specification limits, *ω* the maximum allowable risk of measuring outside these limits and *f*({*M*_*nk*_}) the reportable value. The ATP example can then be formally stated as:

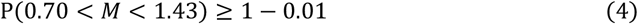

or in layman’s terms: The probability to measure manufactured products (*M*) outside of product specification should not exceed 1 %.

The use of these concrete values is for the sole purpose of making the exposition more comprehensive and illustrative (cf. the discussion section). In practice the % can be much lower, because falling out of specification means facing hard time justifying the result, potentially resulting in incidents and CAPAs – even when the underlying product is compliant.

### Accuracy and precision validation

The accuracy and precision definitions are taken from USP <1033> [2] and rewritten in function of the parameters of the measurement model (Eq. 1). For accuracy this results in:

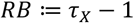

And for precision:

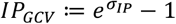

Note that to compute the *RB*, one needs to assume a true or known relative potency value (*μ*_*T*_, cf. Eq. 1) which USP <1033> calls the “ target potency” [2] or “ expected potency” [3]. If the true value is not (assumed) known, then there is no way to estimate *RB*. USP <1033> suggests that for validation purposes target potencies can be constructed “ by dilution of the standard material or [based on] a test sample with known potency”. For *IP* USP <1033> [2] prefers to use the Kirkwood’s [7] GCV formula instead of RSD. Preference of one formula over the other is a matter of taste in this context as they are both transformations of 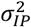 and are used merely for reporting but carrying no deeper meaning or inferential value.

The classical way to validate procedures is by setting global acceptance limits on accuracy (*RB*) and precision (*IP*). To determine them, original USP <1033> [2] uses the process capability index, more specifically the *Ĉ*_*pm*_ variant, which is sometimes called the Taguchi capability index. The latest USP <1033> [3] draft uses an exact solution. It is called “ exact” in this paper because the solution is derived directly from Eq. 3 after substitution of process parameters from the ATP. Both methods result in solutions as global (*RB, IP*) pairs that satisfy the ATP. These solutions are depicted in Fig. 1 as “ Cpm” and “ Exact”, with only distinction being that “ Cpm” is more conservative, imposing unnecessarily strict limits on *RB*.

**Fig. 1:**
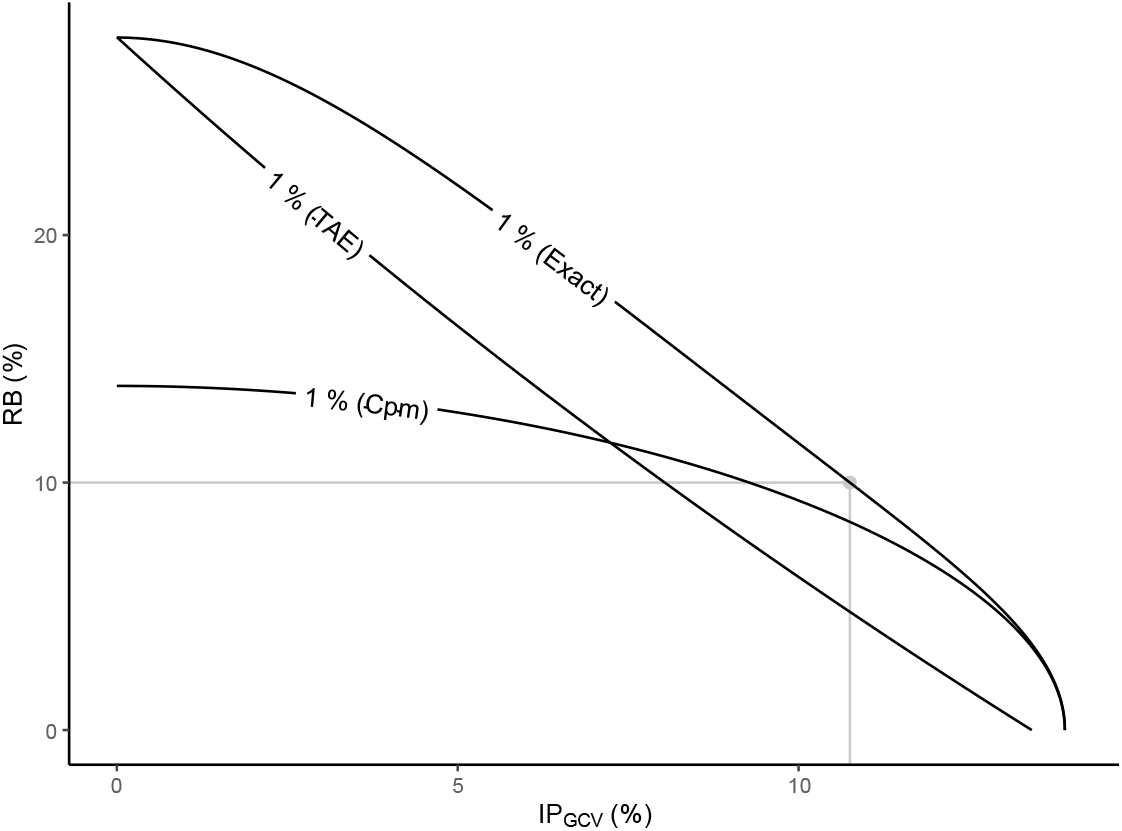
Procedure performance depicted in function of global RB and IP. Curves show the global (RB, IP) pairs that correspond to procedure performance that would result in 1 % out-of-specification measurements under the ATP requirements, computed using _pm_, TAE and the Exact way. The grey rectangle covers all procedures having RB ≤ 10 % and IP_GCV_ ≤ 10.75 %.

It is important to note that the ATP doesn’t imply a single global (*RB, IP*) limit but rather a multitude of limits. Concretely this means that *any* procedure with a set of global (*RB, IP*) parameters that is located below the “ Exact” curve, satisfies the ATP. Unfortunately, in practice only one global (*RB, IP*) pair is taken from the “ Exact” (or even worse the “ Cpm”) curve and is then used as the acceptance limit for the procedure during validation. For example, RB ≤ 10 % and IP_GCV_ ≤ 10.75 %. This pair is depicted by the grey dot in Fig. 1 and all procedures falling within the grey rectangle are deemed acceptable, even though many valid procedures just outside this rectangle (but still below the “ Exact” curve) would also satisfy the ATP, but in practice would be rejected.

A further limitation of this validation methodology is that the acceptance limits are imposed globally across the full working range of the procedure. Yet, as shown in Table I, a procedure’s actual performance is highly dependent on the true value (*μ*_*T*_) that is being measured, meaning that the procedure’s characteristics are local, not global. Imposing global limits can cause issues: For example, the ATP clearly defines the assumed production process under which in our given example it is extremely unlikely to manufacture products close to 2 RP. Yet by setting global (*RB, IP*) acceptance limits one is imposing equal quality standards on the procedure’s performance when measuring at 1 RP as at 2 RP true value. This excessively strict requirement is not warranted by the ATP and will be solved in the correct ATP validation section where the novel validation methodology is presented.

### Total analytical error (TAE) validation

In the previous section on accuracy and precision we established that the Analytical Target Profile (ATP) cannot be satisfied by a single global (*RB, IP*) acceptance limit. This naturally leads us to consider whether the Total Analytical Error (TAE) methodology fares any better.

At its core, the TAE methodology aims to define acceptance limits around the true measured value such that the procedure’s measurements fall within these bounds with a high probability. Formally, this is expressed as:

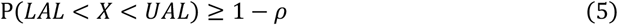

Where *X* is the measured value (cf. Eq. 1), *LAL* and *UAL* the lower and upper acceptance limits for the procedure’s total analytical error (hence the acronym TAE), and *ρ* the maximal risk of observing a result outside this range.

However, even in this initial form, the problem is ill-posed, as Eq. 5 allows for infinitely many valid (*LAL, UAL, ρ*) solutions. To arrive at a unique solution, one typically imposes two additional constraints:

1. In*LAL* and In*UAL* are forced to be symmetric around In*μ*_*T*_. TAE acceptance limit (*TAL*) becomes then: *TAL* ≔ [exp(In*μ*_*T*_ ± In*AL*)].
2. Fixing *ρ* to an arbitrary but intelligible value (i.e. 5 %).

For the reader concerned about “ arbitrary” *ρ*, note that *ρ* and *AL* influence each other: larger *ρ* implies a narrower *AL* and vice versa, but the pair (*AL, ρ*), whatever the initially chosen *ρ* may be, is a unique solution under Eq. 5.

Next, consider the scenario where the analytical procedure is used to measure manufactured products. In this case, the true value *μ*_*T*_ in *X* becomes the random variable *μ*_*P*_*P*, which once substituted in Eq. 5 results in the same form as Eq. 3. This shows that if the production process is perfect (i.e. if *μ*_*P*_*P* always resolves to a fixed RP and hence has 0 variability), then *TAL* and risk (*ρ*) have the same meaning as respectively product specification limits (*L*/*USL*) and probability to measure outside of specification (*ω*), and can be interchanged. This is the approach taken in the latest USP <1033> draft [3] for the TAE validation, which is a gross oversimplification as in practice a production process always has some variability and a translation from product specifications to procedure acceptance limits is required. This translation can be formally stated as follows:

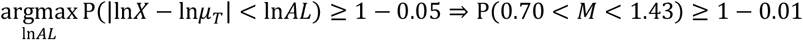

which in layman’s terms says that one is seeking the widest possible *TAL* for the procedure’s total error, such that it guarantees ATP compliance when applied to manufactured products.

The resulting unique solution is In*AL* = 0.247 InRP and *ρ* = 5 %. This solution is depicted as the “ TAE” curve using the corresponding (*RB, IP*) pairs in Fig. 1. The “ TAE” curve shows that any procedure that has a total error within the *TAL* interval at 5 % risk necessarily also complies to the ATP (i.e. the “ TAE” curve is always below or at the “ Exact” curve). Showing that the procedure is within the *TAL* during validation then often results in making an Analytical Error Profile plot as depicted in Fig. 2 from which one can see that the expected 95 % total error interval of the procedure (purple lines) is well within the *TAL* (brown lines).

**Fig. 2:**
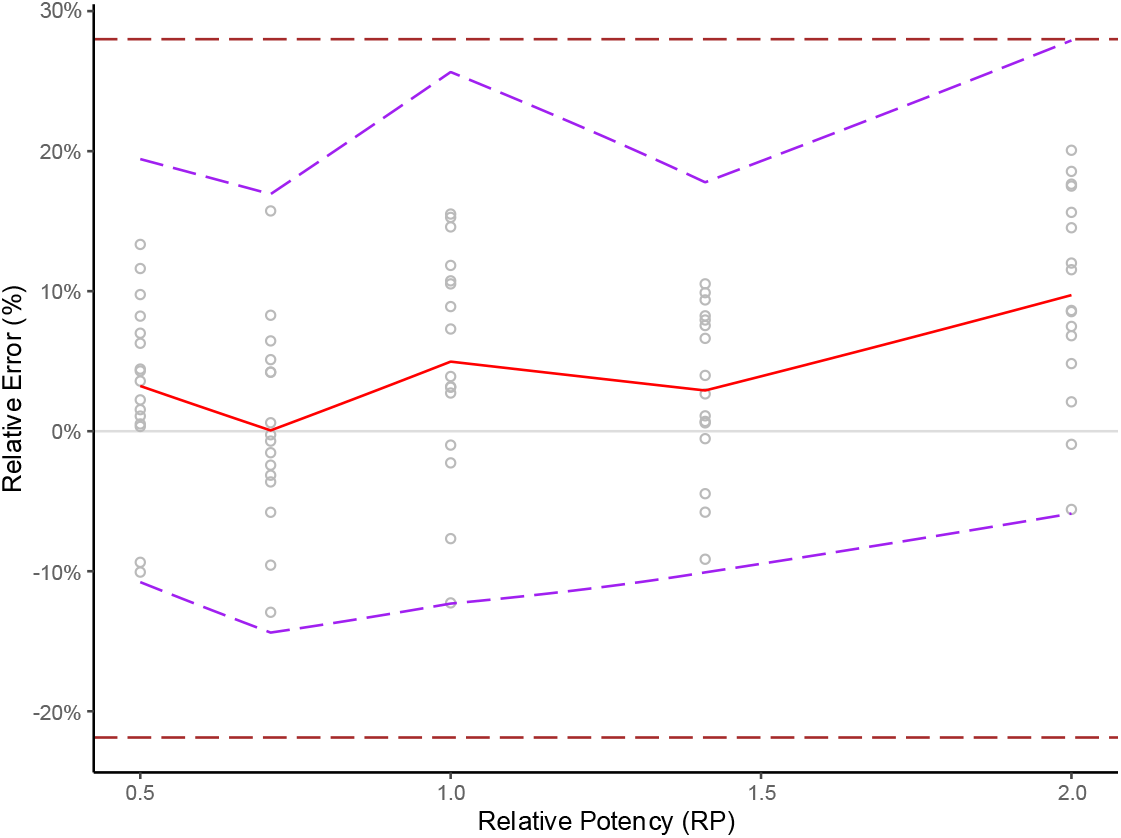
Total Analytical Error profile of the procedure. The experimental measurements (grey circles) are plotted as relative error (%) in function of true relative potency (RP). The red curve is the expected relative bias (RB) and the purple dotted interval is the expected 95 % total error interval of the procedure’s measured values X. The brown dotted horizontal lines are the TAE acceptance limits (TAL).

Compared to accuracy and precision validation methodology, the TAE methodology is more inclusive: It validates any procedure below the “ TAE” curve shown in Fig. 1, which already covers more area than the rectangular regions defined by fixed (*RB, IP*) limits from the “ Exact” or “ Cpm” curves. Yet, limitations remain. The “ TAE” methodology still excludes many valid procedures that are in between the “ TAE” and the “ Exact” curve hence does not fully comply to the ATP requirement. What is more is that the TAE methodology is also based on global limits, just like the accuracy and precision methodology. In other words, a single global *TAL* is applied enforcing equal quality standards across the entire working range.

### Correct (or direct) ATP validation

In previous sections we showed that both accuracy and precision, and TAE and risk methodologies are inadequate to correctly capture the ATP requirements. Both methodologies are susceptible to rejecting valid methods and are based on global acceptance criteria. In this section we propose a methodology for validation that doesn’t try to translate the ATP but uses it directly to validate the procedure. This direct approach avoids the pitfalls of earlier methods and ensures full alignment with local procedure’s performance and the ATP by design.

At the core of the methodology is a direct evaluation of the procedure’s performance with respect to the ATP, as formalized in Eq. 3. The locality of procedure’s performance – which is a direct requirement by Eq. 3 – is modelled with functions In*τ*_*X*_(*μ*_*T*_) and 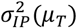 as linear interpolations of the estimated parameters shown in Table I. Other interpolation techniques can also be used. The final solution for the probability to fall out of specification is based on numerical optimization that results in highly accurate estimates of at least 7 significant digits.

While the computation of the correct solution is relatively straightforward, presenting the results in an accessible and actionable way is more challenging. The visualization shown in Fig. 3 represents a practical compromise between complexity and intelligibility, and effectively communicates how the procedure performs relative to the ATP.

**Fig. 3:**
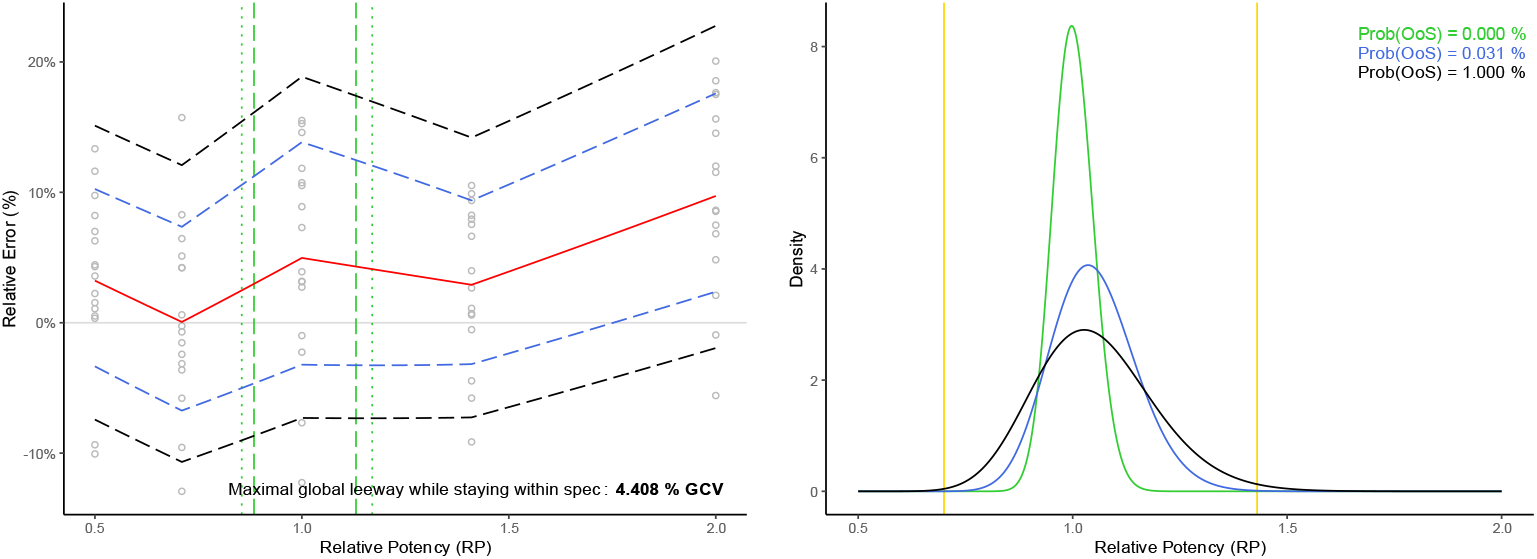
On the left plot the experimental measurements (grey circles) are plotted as relative error (%) in function of true relative potency (RP). The red curve is the expected relative bias (RB) and the blue dotted interval is the added intermediate precision (IP). (These estimates are taken from Table I and interpolated as previously explained.) One can state roughly that the blue dotted interval covers about 68 % of the measurements and gives an idea of the procedure’s measurement performance. The green dashed vertical lines represent the region where the production process is producing 99 % (i.e. 1 – ω) of the products. Hence the procedure’s performance in this region is most important. The dotted green vertical lines represent the region of 99.9 % (i.e. 1 – ω/10) products. The difference between these two green regions gives an idea about the gradient of importance. The density curves in the right plot reflect the performance in routine. The green curve represents the density of true RP of the manufactured products by the production process. The blue curve tells us the density of our reportable values when we measure these products with our procedure (i.e. based on the procedure’s performance summarized with the blue dotted lines). One can see that the measurements would remain well within the boundaries of the product specification (i.e. the yellow bars). The black density has exactly 99 % of its area within the yellow bars, which then translates to the black dotted acceptance interval of the procedure as the maximal addition of global IP to the current performance (blue dotted lines) while still meeting the ATP requirements. Hence the difference between black and blue dotted lines can be interpreted as the maximal global IP that the procedure can incur while still remaining within the ATP specification. The results shown have been validated by simulation for correctness.

The novelties of this correct validation methodology are:

1. *Direct validation against the ATP*: The methodology avoids translation of the ATP into auxiliary acceptance criteria, eliminating the risk of false rejections due to translational errors. The methodology is correct in the sense that it is validating directly against the ATP.
2. *Integration of process knowledge*: The methodology considers performance of the procedure less important in regions where the production process is less likely to produce products (and vice versa). This is consistent with the (assumed) knowledge about the production process embedded in the ATP.
3. *Recognition of local procedure behavior*: Unlike with global acceptance limits, this methodology acknowledges that a procedure may perform well in some parts of the range and less well in others. This local flexibility allows for compensation: strong performance in critical regions can offset weaker performance elsewhere, provided the overall probability of falling out of specification remains within the ATP bounds.

To make this methodology more tangible, Fig. 3 is made interactive in an online demo application [8]. Users can adjust key components – such as the product specifications, out-of-specification risk, the manufacturing distribution, or the procedure’s performance characteristics – and receive immediate feedback on the procedure’s suitability in routine use. This makes it possible to explore trade-offs and optimize the procedure in terms of development effort versus routine impact.

## Discussion

While the focus of this paper’s ATP example has been on validating procedures based on the measured value (i.e. one run and one replicate), extending it to a more complex reportable value and checking its impact based on different formats is straightforward and implemented in the online application [8].

The production process in the ATP has been assumed known and set to 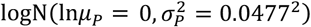 following the example provided in the USP <1033> draft [3]. But within this novel validation framework it could be defined in various ways. For instance, the production process variability could be defined as a “ proportion of the overall [measured] manufacturing variance” [3] (i.e. as proportion of Var[In*M*]). Although this is already suggested in USP <1033> draft [3] the rationale behind the proposed calculation is questionable. There the production process variability 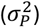 is derived from the reportable value which is based on an (arbitrary) chosen (*RB, IP*) pair from the “ Exact” curve. This is a dangerous practice as production process variability should not depend on an arbitrary chosen *IP* nor on the format of the reportable value, but rather on the 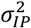, if anything.

The estimates in Fig. 3 are based on point-estimates of procedure’s performance parameters and don’t account for uncertainty due to the limited number of samples in the validation design. To account for this, the ATP could equally well be defined in terms of tolerance limits, e.g. to have 95 % confidence in the validation result.

We deliberately excluded certain aspects of validation, such as dilutional linearity and range determination, which are adequately addressed in USP <1033> [2]. Their treatment remains unchanged under the proposed validation approach.

Importantly, our critique on the accuracy and precision methodology is not aimed at the equivalence testing approach which USP <1033> recommends for assessing whether a procedure meets predefined accuracy and precision limits. Equivalence testing is entirely appropriate provided these acceptance limits are part of the ATP. But when the ATP is specified in terms of product and process characteristics – as in our example – deriving unique accuracy and precision limits is not possible, as shown in earlier sections. The proposed correct methodology validates the procedure directly against the ATP formulation, avoiding the need for surrogate acceptance limits.

We also haven’t assessed the procedure’s Type II errors and its ability to detect defective (i.e., truly out-of-specification) manufactured products simply because this is not within the scope of the validation process as defined in USP <1033> [2]. Nonetheless, evaluating this capability is important from a quality control and risk management perspective. The correct validation methodology presented in this paper enables such an assessment by leveraging the production process knowledge embedded in the ATP as well as the estimated procedure performance. Specifically, one can calculate the probability of correctly identifying products that are truly out of specification, providing quantitative insight into the diagnostic sensitivity of the procedure. This capability has broader implications: these detection probabilities can inform process control strategies, support risk-based decision-making, and align with broader quality-by-design principles advocated in ICH guidelines. As such, we advocate for incorporating this form of analysis alongside traditional validation elements to provide a more complete picture of a procedure’s fitness for purpose.

However, in light of the existing USP <1033> [2] validation requirements, it is also worth reconsidering the common practice of evaluating procedure performance at extreme potency levels, such as 0.5 and 2.0 RP. Under the example ATP used in this paper, the probability that the manufacturing process produces products outside the [0.71, 1.41] RP range is less than 1e-12, i.e. practically zero. If Type II error analysis is not within the validation scope, then testing at extremes such as 0.5 and 2.0 RP becomes unnecessary. Instead, experimental efforts should be concentrated within the feasible production process range, where performance is most relevant and impactful. Broader testing may still be warranted if Type II error assessment is desired (cf. supra), but otherwise, aligning experimental design with the ATP supports more efficient use of resources.

## Conclusion

This paper has demonstrated that validation methodologies based on accuracy and precision, and TAE and risk, fall short when the ATP is defined in terms of product and process specifications, as recommended by USP <1033> [2,3]. Rather than attempting to translate such specifications into surrogate metrics like accuracy, precision and TAE, a more robust approach is to align the validation methodology directly with the ATP itself. To this end, we have proposed a novel, statistically correct validation methodology that avoids the limitations of traditional approaches and ensures full compliance with the ATP. The methodology is also made freely available through an interactive online application at https://apps.rovad.be/usp-1033/. We hope this contribution will stimulate further dialogue on modernizing analytical procedure validation in line with current regulatory expectations.

